# In-silico method for predicting infectious strains of Influenza A virus from its genome and protein sequences

**DOI:** 10.1101/2022.03.20.485066

**Authors:** Trinita Roy, Khushal Sharma, Anjali Dhall, Sumeet Patiyal, Gajendra P. S. Raghava

**Author notes:** **Mailing Address of Authors**, Trinita Roy (TR), Khushal Sharma (KS), Anjali Dhall (AD), Sumeet Patiyal (SP), Gajendra P. S. Raghava (GPSR). Equal Contribution (Trinita Roy, Khushal Sharma, Anjali Dhall). **Corresponding Author**, Prof. Gajendra P. S. Raghava, Head and Professor, Department of Computational Biology, Indraprastha Institute of Information Technology, Delhi, Okhla Industrial Estate, Phase III, New Delhi, India – 110020 Office: A-302 (R&D Block), Website: http://webs.iiitd.edu.in/raghava/. **Author’s Biography** 1. Trinita Roy is currently pursuing MTech in Computational Biology from Department of Computational Biology, Indraprastha Institute of Information Technology, New Delhi, India. 2. Khushal Sharma is currently pursuing MTech in Computational Biology from Department of Computational Biology, Indraprastha Institute of Information Technology, New Delhi, India. 3. Anjali Dhall is currently working as Ph.D. in Computational Biology from Department of Computational Biology, Indraprastha Institute of Information Technology, New Delhi, India. 4. Sumeet Patiyal is currently working as Ph.D. in Computational Biology from Department of Computational Biology, Indraprastha Institute of Information Technology, New Delhi, India. 5. Gajendra P. S. Raghava is currently working as Professor and Head of Department of Computational Biology, Indraprastha Institute of Information Technology, New Delhi, India.

## Abstract

Influenza A is a contagious viral disease responsible for four pandemics in the past and a major public health concern. Being zoonotic in nature, the virus can cross the species barrier and transmit from wild aquatic bird reservoirs to humans via intermediate hosts. Virus gradually undergoes host adaptive mutations in their genome and proteins, resulting in different strain s/vari ants which might spread virus from avians/mammals to humans. In this study, we have developed an in-silico models to identify infectious strains of Influenza A virus, which has the potential of getting transmitted to humans, from its whole genome/proteins. Firstly, machine learning based models were developed for predicting infectious strains using composition of 15 proteins of virus. Random Forest based model of protein Hemagglutinin, achieved maximum AUC 0.98 on validation data using dipeptide composition. Secondly, we obtained maximum AUC of 0.99 on validation dataset using one-hot-encoding features of each protein of virus. Thirdly, models build on DNA composition of whole genome of Influenza A, achieved maximum AUC 0.98 on validation dataset. Finally, a web-based service, named “FluSPred”(https://webs.iiitd.edu.in/raghava/fluspred/) has been developed which incorporate best 16 models (15 proteins and one based on genome) for prediction of infectious strains of virus. In addition, we provided standalone software for the prediction and scanning of infectious strains at large-scale (e.g., metagenomics) from genomic/proteomic data. We anticipate this tool will help researchers in prioritize high-risk viral strains of novel influenza virus possesses the capability to spread human to human, thereby being useful for pandemic preparedness and disease surveillance.

**Key Points:**

- Influenza A is a contagious viral disease responsible for four pandemics.
- Virus can cross species barrier and infect human beings.
- In silico models developed for predicting human infectious strains of virus.
- Models developed were build using 15 proteins and whole genome datasets.
- Webserver and standalone package for predicting and scanning of high-risk viral strains.

## Introduction

Influenza is a contagious viral disease, having a zoonotic origin, where a sudden rise in body temperature, headache, myalgia, lethargy, and dry cough [1] are some of the symptoms of the infection. Severe cases ranging from influenza-induced pneumonia [2], encephalitis, myocarditis [3], to death within a few hours, can have fatal outcomes. Patients with chronic heart or lung disease [4], immune disorders, diabetes, have an increased risk of influenza infection [5–7]. Influenza A virus occurs naturally among wild aquatic birds like geese, swans, waterfowl and it is responsible for causing avian influenza in birds, including domestic poultry [8]. However, virues jump species barriers to affect animals and humans sporadically [9]. As shown in Figure 1, different virus strains can affect humans and cause epidemics as the strains may have the potential to overcome species barriers and infect humans. Influenza A virus may further get adapted in such a way that it could spread human-to-human, leading to a potential pandemic, which is a cause of great concern [10, 11].

**Figure 1:**
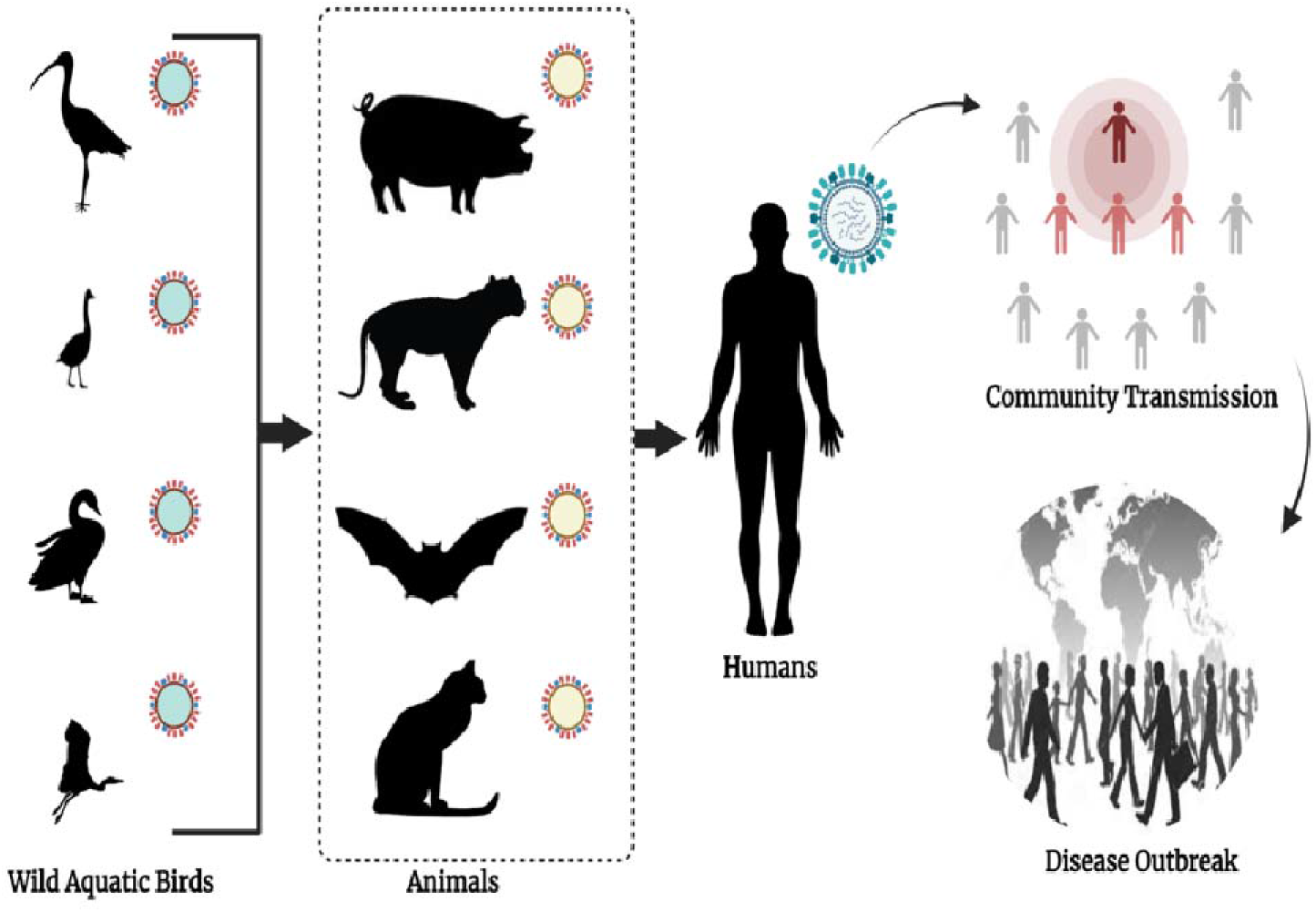
Schematic representation of the zoonotic spread of Influenza A from wild aquatic birds to animals by jumping the species barrier and eventually infect humans which can lead to human-to-human, community transmission to disease outbreaks like epidemics and pandemics.

**Figure 2:**
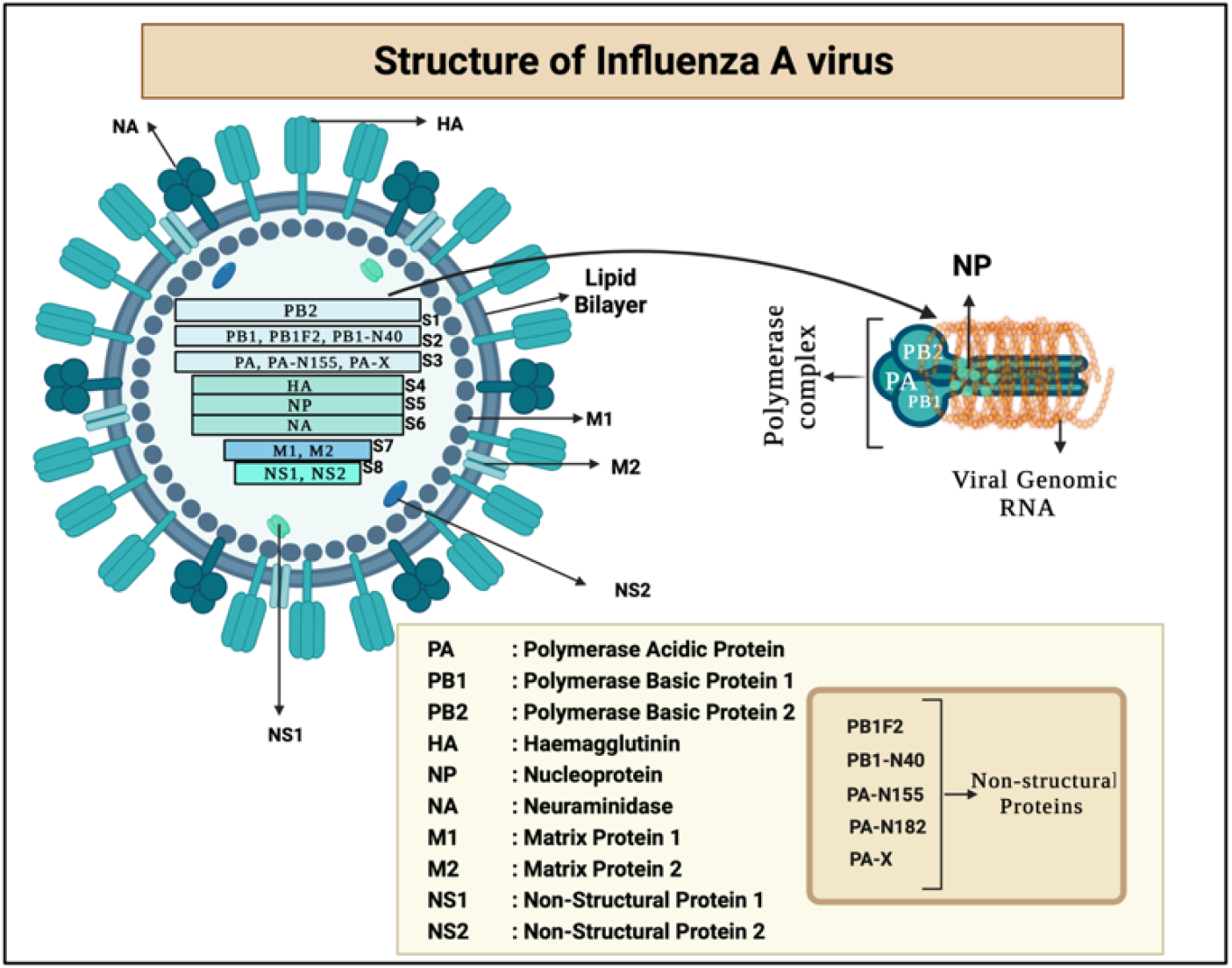
Schematic representation of Influenza A Virus with a total of eight segments whereas, (PA, PB1, and PB2) are the subunits of RNA-dependent RNA Polymerase proteins, HA and NA are surface glycoproteins, (NP) RNA binding protein, (M1) bifunctional membrane/RNA-binding protein, (M2) ion-channel protein, (PB1F2, PA-N155, PA-N182, and PA-X) are non-structural proteins.

The pandemic influenza strains arise from reassortment of gene segments, where there is major antigenic change occurs, i.e., a phenomenon known as antigenic shift [12]. The non-human viruses undergo changes so that they become capable of infecting humans and spreading efficiently and sustainably [13]. In other words, mutations in amino acid residues, especially haemagglutinin, can alter the receptor-binding site and specificity, altering receptor preference and spread at large scale in various countries [14]. In past, Influenza A was responsible for the outbreak of four pandemics. One of the most devastating pandemics, with a high mortality rate among children below 5 years of age and 20-40 age group, was caused by the H1N1 subtype in 1918. The virus had an avian origin and resulted in 50 million estimated deaths worldwide [15]. The H2N2 subtype emerged in East Asia from an avian source in 1957 causing approximately 1.1 million deaths worldwide [16].

In 1968, the H3N2 subtype was first noted in the USA and had over 1 million deaths globally. This virus later started circulating as the seasonal flu [17]. In 2009 (H1N1) pdm09 was first noted in the USA and leads to 151,700-575,400 deaths worldwide in the first year of infection, then it started circulating as seasonal flu [18]. Host adaptive mutations in the genome and proteins are the major reasons behind the transmissibility of the viral strains from the avian and mammalian reservoirs to human hosts [19]. In the recent past, several methods have been developed for the identification of host tropism of Influenza A virus [20–23] using protein and genome sequences. These studies show that protein and genome sequences are very efficient for determining the host tropism of zoonotic pathogens and changes take place at the molecular level for the virus being capable of crossing the species barrier.

In this study, we have made a systematic attempt to develop computational models for the prediction of infectious strains of the influenza A virus. The aim of this study is to monitor infectious strains in non-human reservoirs to control the spread of strain from non-human to human hosts. We compiled 15 influenza A virus proteins and genome sequences from IRD and VIDHOP resources respectively. Models using different machine learning techniques were developed where features of proteins and genome were computed using Pfeature and Nfeature. In order to serve the scientific community, we provide a freely accessible web-server, and a standalone package named “FluSPred”, for the prediction of infectious influenza A strains with the help of either of the proteins or the genome sequence using the best prediction models.

## Material and Methods

The complete architecture of the study is illustrated in Figure 3. The description of each step is given below.

**Figure 3:**
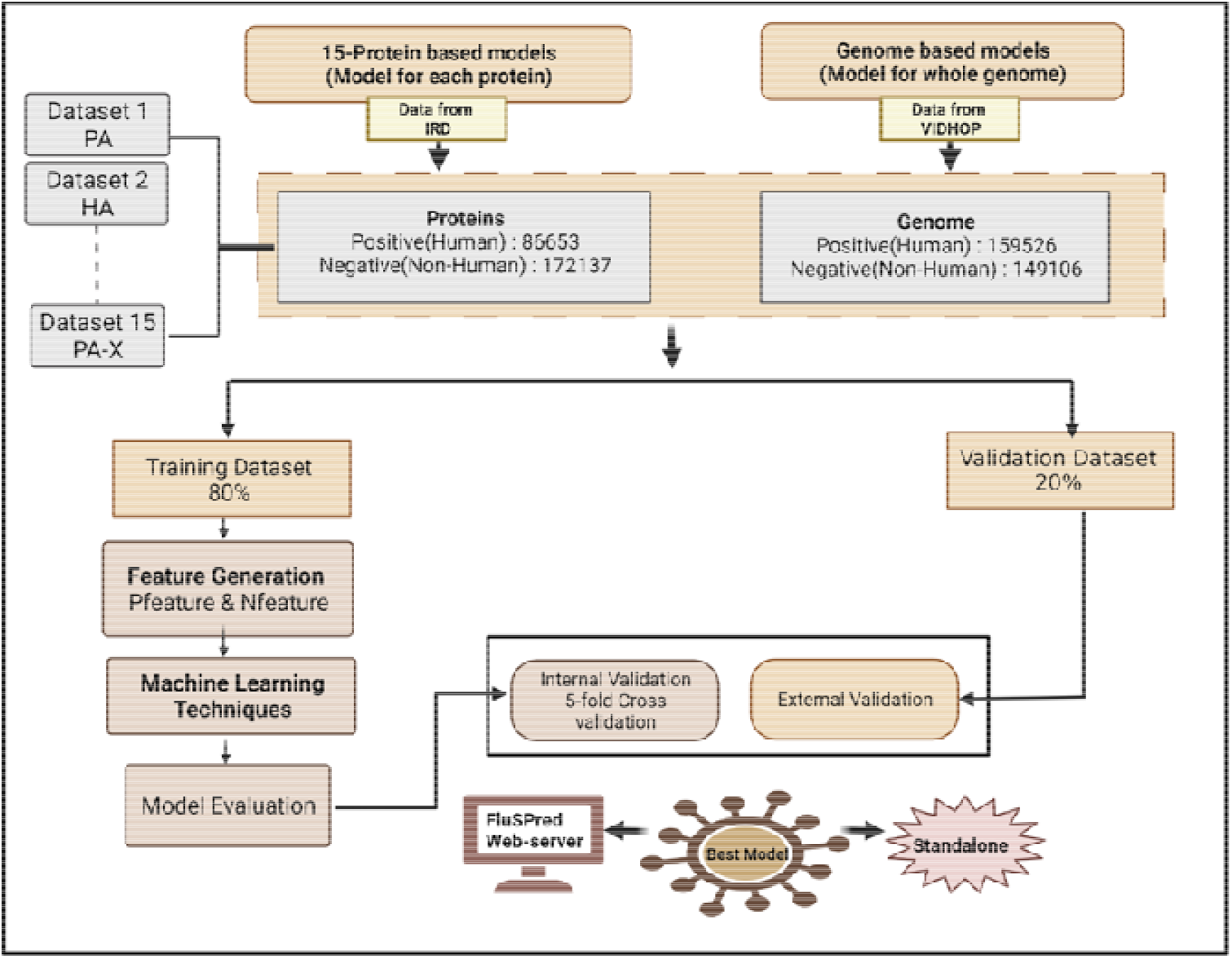
The flowchart explains the work and design of FluSPred, including dataset collection, feature generation, and machine learning techniques.

### Datasets for protein-based models

We obtained a total of 985,720 protein sequences from the Influenza Research Database (IRD) database (www.fludb.org) corresponding to 15 proteins of the Influenza A virus. These proteins were used to create 15 datasets (one dataset for one type of protein), where each dataset has sequences of the viral proteins from human host and non-human host (avian/mammalian host). In order to avoid biases, we removed the redundant and incomplete sequences (See Table 1). In this study, human infectious strains and associated sequences were considered positive datasets, and non-human (avian and other mammalian) was taken as a negative dataset. After preliminary data cleaning, we get 258790 unique protein sequences out of which, 86653 were positive (i.e. viral protein sequences derived from human hosts) and 172137 were negative sequences (i.e., viral protein sequences derived from non-human hosts). The comprehensive distribution of the dataset for different Influenza A proteins is provided in the table below.

**Table 1:** Distribution of positive and negative dataset for the Influenza A protein sequences

| Protein Name | Positive Dataset<br>(Human) | Negative Dataset<br>(Non-Human) | Total Dataset |
| --- | --- | --- | --- |
| Polymerase Acidic Protein(PA) | 7722 | 14651 | 22373 |
| Polymerase Basic Protein 1(PB1) | 6011 | 12407 | 18418 |
| Polymerase Basic Protein 2(PB2) | 7961 | 14165 | 22126 |
| Hemagglutinin(HA) | 17999 | 27350 | 45349 |
| Nucleoprotein(NP) | 3716 | 9031 | 12747 |
| Neuraminidase(NA) | 13486 | 20315 | 33801 |
| Matrix Protein 1(M1) | 1309 | 2937 | 4246 |
| Matrix Protein 2(M2) | 1706 | 4533 | 6239 |
| Non-Structural Protein 1(NS1) | 5577 | 10414 | 15991 |
| Non-Structural Protein 2(NS2) | 1376 | 4158 | 5534 |
| PB1-F2 | 2298 | 9441 | 11739 |
| PB1-N40 | 5028 | 10893 | 15921 |
| PA-N155 | 4687 | 10716 | 15403 |
| PA-N182 | 4422 | 9739 | 14161 |
| PA-X | 3355 | 11387 | 14742 |

### Dataset for genome-based models

Influenza A genomic sequences were obtained from the VIDHOP method [24]. A total of 312617 genome sequences were available in the dataset in which 159526 humans and 153091 nonhuman genomes were selected for this study. In order to maintain the homogeneity in protein and genome datasets, here we considered only avian, mammalian, and human. Viral genome sequences derived from human hosts were taken as positive and viral genome sequences derived from non-human hosts were taken as the negative dataset. Apart from this, the duplicate sequences were removed, and finally, a total of 308632 sequences were considered, with 159526 positive sequences and 149106 negative sequences.

In order to train and evaluate our models, we perform both internal and external validation. Thus, the final dataset was divided into training and validation data, where 80% data was used for internal and 20% data was used for external validation [25–28].

### Feature Generation

#### Composition-based features

In order to compute the amino acid composition (AAC) and dipeptide composition (DPC) features of protein sequences, we have used the Pfeature standalone package [29]. In case of AAC, a feature vector of the length of 20, and for the DPC, a vector length of 400 features was computed. Furthermore, to generate the nucleotide-based features for the genome sequence data, we have used Nfeature standalone package [30]. Here, we computed composition-based features such as the Composition of DNA sequence for all K-Mers (CDK) and Reverse complement of DNA for all K-mers (RDK). CDK is a feature extraction method that calculates the composition of all 4 nucleotides. Here K could be 1, 2, or 3, generating a feature vector of length 4, 16, and 64 respectively. Likewise, RDK is a feature extraction method that calculates reverse complement K-Mer composition of nucleotides where we used K as 1,2 and 3 having feature vector sizes of 2,10, and 32 respectively.

#### One-hot Encoding

In addition, we have used one hot encoding approach for feature extraction, where the length of the sequences needed to be equal. In order to generate equal length vectors, we fix the length of all the sequences to 0.95 quantiles, where the sequences having shorter than that were extended by normal repeats, and the longer sequences were truncated until it reaches 0.95 quantiles of all sequences mark [24]. Once the lengths of the sequences are equalized, the sequences are encoded numerically by the one-hot encoding method. Here, each of the amino acid residues of the entire sequence is converted to a vector size of length of the sequence x 20, where 20 is the canonical protein residues, where the respective residue is given the value 1 in its position and the rest is given the value 0. This generates a sparse matrix, for instance, Alanine (A) can be represented by a 20-dimensional vector 10000000000000000000. Similarly, Cysteine(C) can be represented as a 20-dimensional vector 001000000000000000000.

#### Machine Learning Models

In this study, several machine learning algorithms have been implemented for developing binary classification models that include Random Forest (RF), K-Nearest Neighbours (KNN), Decision Tree (DT), Support Vector Machine (SVM), XGBoost (XGB), Gaussian Naive Bayes (GNB). KNN is an instance-based classification method that classifies based on the vote of the nearest neighbor data points. It only stores the instances of the training variables and classification is determined from the majority vote of the nearest neighbor of each data point. Whereas, RF, is an ensemble-based method for classification, while training, the RF classifier fits numerous DTs to predict the response variable as an individual tree. Averaging DTs improves the prediction accuracy and control on overfitting of the models. However, DT is a non-parametric-based supervised learning algorithm that predicts the response variable by learning the decision rules from the data features. GBM algorithm is a probabilistic classification approach based on Bayes theorem which assumes that the continuous variables of each class follow the normal or Gaussian distribution XGB is an ensemble method that implements the gradient boosted decision trees designed for speed and performance. It uses an iterative approach and provides a wrapper class to treat models like classifiers or regressors. For each of the 15 proteins, SVM, RF, and KNN were used on AAC, DPC, One Hot Encoding features. On the other side, DT, KNN, XGB, GNB, and RF-based models were generated for genome sequences using CDK and RDK features. We fine-tuned the model on the training data by taking different thresholds for probabilities as 0.8,0.5, 0.4, 0.2, 0.1, and we found that 0.5 works best for all the models. These classification techniques were implemented using python-library sci-kit learn [31].

#### Feature Selection

Identifying the critical set of features for the machine learning models is one of the significant challenges in this study. For one hot encoding feature generation technique, each residue of the protein sequences was represented as a 20-dimensional vector (sequence length x 20 dimensions) where the resultant matrix became sparse. Therefore, we needed dimensionality reduction where we could reduce and choose the final number of components which aids to the faster execution of the program as well as reduces the noise in the data. We have used the Truncated Singular Value Decomposition (tSVD) method for one-hot encoded feature selection to improve computational efficiency. The highly dimensional matrix was then reduced to 100 principal components by calculating the explained variance [32].

#### Cross-Validation

The dataset was split into an 80:20 ratio, where 80 percent data consisted of training data and the rest 20 percent was validation data. The training data was further divided into training and testing datasets during the 5-fold cross-validation process and the mean of the results for each fold of the cross-fold validation was noted down. In the 5-fold cross-validation process, the entire training data gets divided into five equivalent folds and then four folds were used for training and the 5th fold was used for testing. The whole procedure gets iterated five times where every fold gets a chance for being used as testing data. This is a standard procedure used in many studies [33–37].

#### Evaluation Parameters

In the current study, we have used the standard performance evaluation parameters such as accuracy, sensitivity, specificity, precision, recall, Matthews Correlation Coefficient (MCC), and Area Under the Curve (AUC) were computed. Whereas, accuracy, sensitivity, and specificity are threshold-dependent parameters and AUC is threshold independent parameter that is plotted by sensitivity vs 1-specificity. The parameters are computed by:

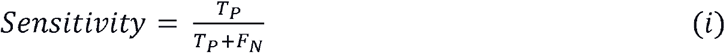

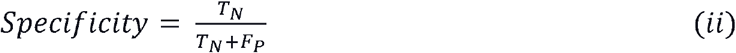

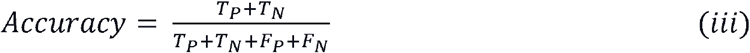

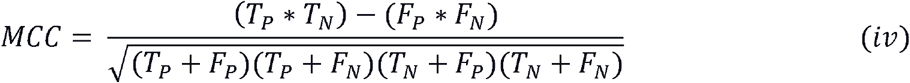

Where, T_P_, T_N_, F_P_ and F_N_ stand for true positive, true negative, false positive and false negative, respectively.

## Results

### Composition Based Analysis

The average amino acid composition of positive and negative sequences for 15 proteins were computed to determine the changes in the average compositions of proteins. In the Supplementary Figure S1, we have provided the amino-acid based compositional analysis for all the protein sequences. Here, we are showing an example of only HA and NA protein, since they play a significant role in the zoonosis as their variations are responsible for the different subtypes of Influenza A [38, 39]. As shown in Figure 4, the average composition of Alanine (A), Isoleucine (I), and Lysine(K), is comparatively higher in the positive dataset, where, the Glutamic Acid (E), Glycine (G), Leucine (L), and Methionine (M) shows higher composition in negative dataset for HA protein. Likewise, for NA protein, Alanine(A), Glutamic Acid(E), Histidine(H), and Asparagine(N) have a higher average composition in the positive dataset whereas Glycine(G) and Tyrosine(Y) are higher in the negative dataset.

**Figure 4:**
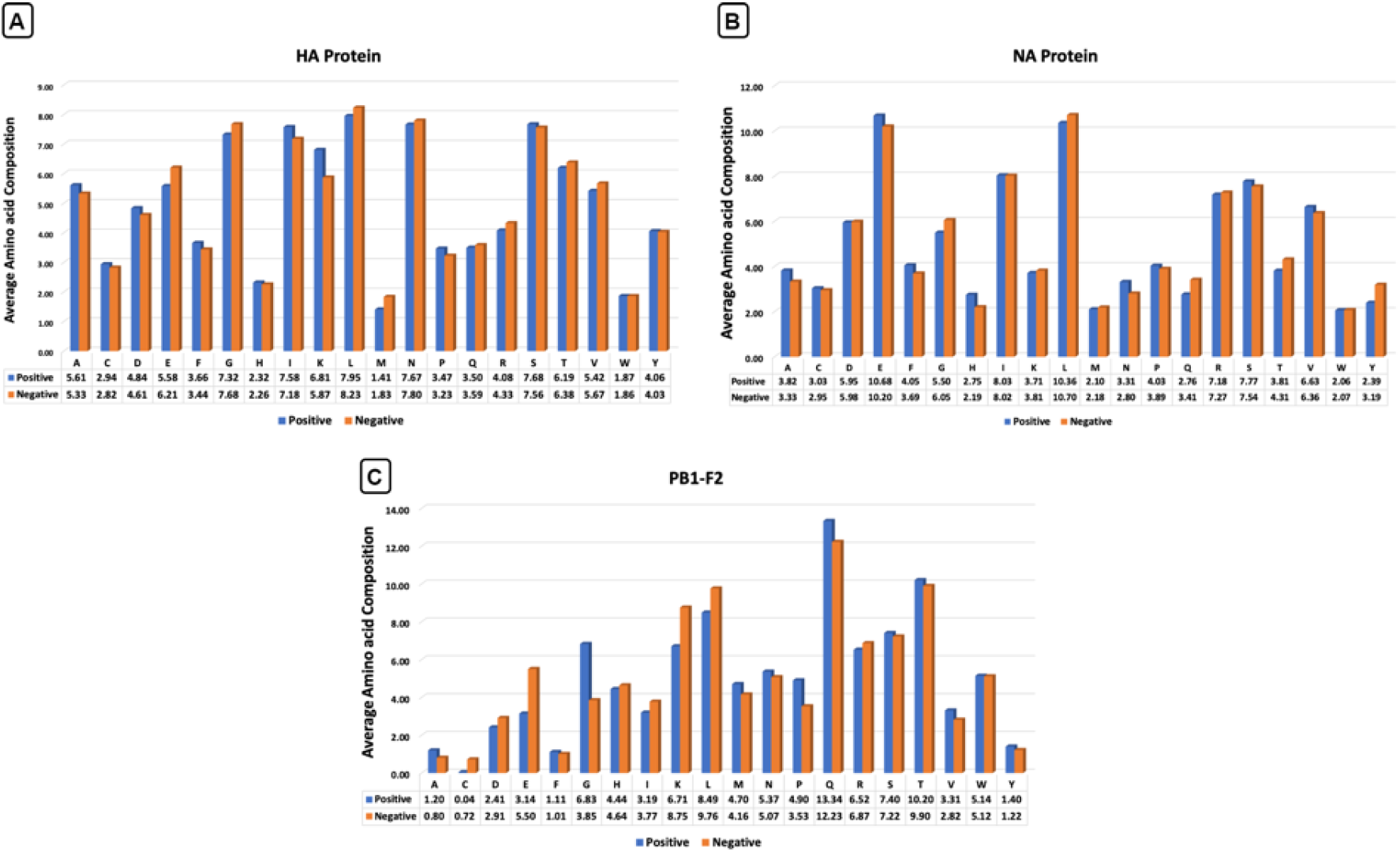
Average amino acid composition of (A) HA protein and (B) NA protein (C) PB1F2 protein

Similarly, we have computed the compositional analysis for the rest of the proteins, as shown in Supplementary Figure S1. Each protein shows a difference in the average composition between the positive and negative datasets. In PB1-F2, for example, the average composition of Glycine(G), Proline (P), and Glutamine (Q) is remarkably high. Consequently, Cysteine(C), Glutamic Acid(E), Lysine (K), and Leucine(L) compositions are significantly higher in the negative dataset as shown in Figure 4. This shows that there is a difference in the compositions in the amino acids which play a role in the virus being capable of crossing the species barrier to infect humans.

### Motif based analysis

In order to perceive the different motifs occurring exclusively in either of the positive or negative datasets/the positive datasets of each of the proteins and the genome, we used MERCI software [40]. It is a tool that can identify and return top K motifs that are most frequent in the positive sequences and absent in the negative sequences. The top 10 motifs of each of the proteins which are exclusively present in the positive and negative datasets are enclosed in Supplementary Table S1.

### Performance of Machine Learning Models

#### Prediction based on protein datasets

Machine learning based models have been developed for all the Influenza A proteins for the prediction of the human infectious influenza virus strains using different types of sequence-based features such as AAC, DPC and One Hot Encoding. The models built using AAC features achieves maximum AUC of 0.973 on validation data with an accuracy of 97.5% for HA protein, as shown in Table 2. Similarly, PB1-F2 prediction models show the highest accuracy of 98.8% on training data as well as validation dataset. Whereas, the other models achieved comparable performances on both training and validation datasets as provided in the table below. The complete results were provided in Supplementary Table S2.

**Table 2:** The performance of Random Forest-based models developed using AAC features for training and validation datasets.

|  | TRAINING |  |  |  |  | VALIDATION |  |  |  |  |
| --- | --- | --- | --- | --- | --- | --- | --- | --- | --- | --- |
| *Proteins* | *Sensitivity* | *Specificity* | *Accuracy* | *AUC* | *MCC* | *Sensitivity* | *Specificity* | *Accuracy* | *AUC* | *MCC* |
| PA | 0.95 | 0.98 | 0.97 | 0.97 | 0.94 | 0.95 | 0.96 | 0.96 | 0.96 | 0.91 |
| HA | 0.96 | 0.99 | 0.98 | 0.97 | 0.95 | 0.98 | 0.97 | 0.97 | 0.97 | 0.94 |
| M1 | 0.95 | 0.94 | 0.94 | 0.94 | 0.86 | 0.73 | 0.95 | 0.88 | 0.84 | 0.71 |
| M2 | 0.94 | 0.97 | 0.97 | 0.96 | 0.91 | 0.83 | 0.96 | 0.93 | 0.90 | 0.81 |
| NA | 0.96 | 0.98 | 0.98 | 0.97 | 0.95 | 0.97 | 0.97 | 0.97 | 0.97 | 0.94 |
| NP | 0.96 | 0.98 | 0.98 | 0.97 | 0.94 | 0.95 | 0.98 | 0.97 | 0.97 | 0.94 |
| NS1 | 0.95 | 0.98 | 0.97 | 0.97 | 0.94 | 0.96 | 0.97 | 0.96 | 0.96 | 0.92 |
| NS2 | 0.93 | 0.98 | 0.97 | 0.96 | 0.93 | 0.89 | 0.96 | 0.94 | 0.93 | 0.85 |
| PB1 | 0.94 | 0.98 | 0.96 | 0.96 | 0.92 | 0.92 | 0.96 | 0.94 | 0.94 | 0.88 |
| PB2 | 0.95 | 0.98 | 0.97 | 0.96 | 0.93 | 0.95 | 0.96 | 0.96 | 0.96 | 0.91 |
| PB1-F2 | 0.99 | 0.99 | 0.99 | 0.99 | 0.96 | 0.92 | 0.99 | 0.98 | 0.96 | 0.94 |
| PB1-N40 | 0.94 | 0.97 | 0.96 | 0.96 | 0.91 | 0.91 | 0.97 | 0.95 | 0.94 | 0.88 |
| PA-N155 | 0.94 | 0.98 | 0.97 | 0.96 | 0.92 | 0.92 | 0.97 | 0.95 | 0.94 | 0.89 |
| PA-N182 | 0.95 | 0.98 | 0.97 | 0.97 | 0.93 | 0.94 | 0.96 | 0.95 | 0.95 | 0.90 |
| PA-X | 0.95 | 0.98 | 0.97 | 0.96 | 0.92 | 0.86 | 0.98 | 0.95 | 0.92 | 0.86 |

Further, we developed various models using the dipeptide composition-based features, where the random forest classifier shows higher efficiency compared to the rest of the models using the same features Supplementary Table S3. Here, the HA protein model showed the best results, in terms of accuracy 97.7% on training as well as validation dataset. Whereas, NP based models give maximum performance with an accuracy of 98.4% and AUC 0.99 on the validation dataset. The complete results of other datasets are shown in Table 3.

**Table 3:** The performance of Random Forest-based models developed using DPC features for training and validation datasets.

|  | TRAINING |  |  |  |  | VALIDATION |  |  |  |  |
| --- | --- | --- | --- | --- | --- | --- | --- | --- | --- | --- |
| *Proteins* | *Sensitivity* | *Specificity* | *Accuracy* | *AUC* | *MCC* | *Sensitivity* | *Specificity* | *Accuracy* | *AUC* | *MCC* |
| PA | 0.96 | 0.99 | 0.97 | 0.97 | 0.94 | 0.97 | 0.97 | 0.97 | 0.97 | 0.93 |
| HA | 0.96 | 0.99 | 0.98 | 0.98 | 0.95 | 0.98 | 0.97 | 0.98 | 0.98 | 0.95 |
| M1 | 0.96 | 0.97 | 0.96 | 0.96 | 0.91 | 0.82 | 0.97 | 0.92 | 0.89 | 0.81 |
| M2 | 0.95 | 0.97 | 0.97 | 0.96 | 0.92 | 0.89 | 0.98 | 0.95 | 0.93 | 0.88 |
| NA | 0.96 | 0.99 | 0.98 | 0.98 | 0.96 | 0.98 | 0.97 | 0.97 | 0.97 | 0.94 |
| NP | 0.98 | 0.98 | 0.98 | 0.98 | 0.96 | 0.96 | 0.99 | 0.98 | 0.98 | 0.96 |
| NS1 | 0.96 | 0.98 | 0.98 | 0.97 | 0.95 | 0.96 | 0.97 | 0.96 | 0.96 | 0.92 |
| NS2 | 0.99 | 0.99 | 0.99 | 0.99 | 0.99 | 0.90 | 0.96 | 0.91 | 0.86 | 0.76 |
| PB1 | 0.96 | 0.99 | 0.98 | 0.97 | 0.95 | 0.94 | 0.97 | 0.96 | 0.96 | 0.91 |
| PB2 | 0.96 | 0.99 | 0.98 | 0.98 | 0.96 | 0.97 | 0.97 | 0.97 | 0.97 | 0.93 |
| PB1-F2 | 0.99 | 0.99 | 0.99 | 0.99 | 0.99 | 0.87 | 0.99 | 0.99 | 0.83 | 0.75 |
| PB1-N40 | 0.96 | 0.98 | 0.98 | 0.97 | 0.94 | 0.95 | 0.97 | 0.97 | 0.96 | 0.92 |
| PA-N155 | 0.94 | 0.99 | 0.97 | 0.96 | 0.93 | 0.95 | 0.97 | 0.97 | 0.96 | 0.92 |
| PA-N182 | 0.95 | 0.98 | 0.98 | 0.97 | 0.94 | 0.95 | 0.97 | 0.97 | 0.96 | 0.92 |
| PA-X | 0.95 | 0.98 | 0.98 | 0.97 | 0.93 | 0.94 | 0.98 | 0.97 | 0.96 | 0.92 |

In addition, we have used binary or one-hot encoding features for the classification of human and non-human sequences. Here, we also observed that HA protein-based models achieved a maximum accuracy (99.9% and 98.3%), and AUC (0.980 and 0.988) on training and validation data, respectively. Moreover, it shows the highest MCC of 0.966 on the validation data as depicted in Table 4. The comprehensive results are provided in Table 4 and the complete results are given in Supplementary Table S4.

**Table 4:** The performance of Random Forest-based models developed using one hot encoding feature for training and validation datasets.

| Proteins | TRAINING |  |  |  |  | VALIDATION |  |  |  |  |
| --- | --- | --- | --- | --- | --- | --- | --- | --- | --- | --- |
|  | Sensitivity | Specificity | Accuracy | AUC | MCC | Sensitivity | Specificity | Accuracy | AUC | MCC |
| PA | 0.97 | 0.99 | 0.98 | 0.98 | 0.96 | 0.96 | 0.97 | 0.97 | 0.97 | 0.93 |
| HA | 0.99 | 0.99 | 0.99 | 0.99 | 0.98 | 0.98 | 0.99 | 0.98 | 0.98 | 0.97 |
| M1 | 0.99 | 0.98 | 0.98 | 0.98 | 0.96 | 0.89 | 0.94 | 0.92 | 0.91 | 0.82 |
| M2 | 0.99 | 0.99 | 0.99 | 0.99 | 0.97 | 0.85 | 0.98 | 0.94 | 0.91 | 0.86 |
| NA | 0.97 | 0.99 | 0.98 | 0.98 | 0.96 | 0.97 | 0.98 | 0.97 | 0.97 | 0.94 |
| NP | 0.99 | 0.98 | 0.99 | 0.99 | 0.97 | 0.93 | 0.98 | 0.97 | 0.96 | 0.92 |
| NS1 | 0.99 | 0.99 | 0.99 | 0.99 | 0.97 | 0.93 | 0.98 | 0.97 | 0.96 | 0.92 |
| NS2 | 0.98 | 0.98 | 0.98 | 0.98 | 0.95 | 0.88 | 0.97 | 0.94 | 0.92 | 0.85 |
| PB1 | 0.97 | 0.99 | 0.98 | 0.98 | 0.96 | 0.94 | 0.97 | 0.96 | 0.96 | 0.91 |
| PB2 | 0.98 | 0.99 | 0.98 | 0.98 | 0.96 | 0.95 | 0.98 | 0.97 | 0.97 | 0.93 |
| PB1-F2 | 0.98 | 0.98 | 0.99 | 0.99 | 0.96 | 0.90 | 0.99 | 0.98 | 0.95 | 0.93 |
| PB1-N40 | 0.97 | 0.98 | 0.98 | 0.97 | 0.95 | 0.95 | 0.97 | 0.96 | 0.96 | 0.91 |
| PA-N155 | 0.97 | 0.99 | 0.98 | 0.98 | 0.96 | 0.93 | 0.98 | 0.96 | 0.95 | 0.91 |
| PA-N182 | 0.82 | 0.94 | 0.90 | 0.88 | 0.77 | 0.85 | 0.91 | 0.89 | 0.88 | 0.75 |
| PA-X | 0.98 | 0.98 | 0.98 | 0.98 | 0.94 | 0.85 | 0.98 | 0.95 | 0.92 | 0.86 |

From the results, we observed that it is evident that although HA and NA are prevalently known to play a role in infection by giving rise to new subtypes or variants, they are not the only ones. The other proteins also act as human infection-causing strains. There is the scope of further research on sequence analysis or molecular studies on these proteins to identify their roles. In addition, we found that proteins like NS1, NS2 or M1, M2 which get encoded from the same segment, respectively, give different prediction results. This shows that even though they are encoded by the same segments, they are functionally different. This may be attributed to both proteins responding individually to structural constraints or selective pressures since a mutation in the respective segment can affect either protein separately [41].

#### Prediction based on genome datasets

In order to develop the genome-based classification models, we have used various machine learning techniques such as RF, KNN, SVM, DT, NB, and XGB. Here, we have computed RDK and CDK based compositional features for the nucleotide sequences, which were used to develop prediction models for finding the human/non-human infectious strains. Likewise, we have generated CDK based features, where the random forest achieved the highest accuracy (97.6%) and AUC of 0.97 on the training data and 98% accuracy and AUC of 0.98 on the validation data, as shown in the table below (Table 5). Here, KNN showed almost similar results with an accuracy of 97.6% and AUC of 0.975 on training data and 97.7% accuracy, and AUC of 0.977 on the validation dataset. The rest of the results for CDK are provided in Supplementary Table S5.

**Table 5:** Performance of machine learning based models developed using CDK (K = 3) features on training and validation dataset.

|  | TRAINING |  |  |  |  | VALIDATION |  |  |  |  |
| --- | --- | --- | --- | --- | --- | --- | --- | --- | --- | --- |
| *Classifier* | *Sensitivity* | *Specificity* | *Accuracy* | *AUC* | *MCC* | *Sensitivity* | *Specificity* | *Accuracy* | *AUC* | *MCC* |
| RF | 0.98 | 0.98 | 0.98 | 0.98 | 0.96 | 0.98 | 0.98 | 0.98 | 0.98 | 0.96 |
| KNN | 0.99 | 0.96 | 0.98 | 0.98 | 0.95 | 0.99 | 0.97 | 0.98 | 0.98 | 0.96 |
| SVM | 0.85 | 0.78 | 0.82 | 0.82 | 0.64 | 0.86 | 0.78 | 0.82 | 0.82 | 0.64 |
| DT | 0.96 | 0.96 | 0.96 | 0.96 | 0.91 | 0.96 | 0.95 | 0.96 | 0.50 | 0.00 |
| NB | 0.74 | 0.61 | 0.68 | 0.68 | 0.36 | 0.74 | 0.60 | 0.68 | 0.67 | 0.35 |
| XGB | 0.96 | 0.94 | 0.95 | 0.95 | 0.89 | 0.96 | 0.94 | 0.95 | 0.95 | 0.89 |

On the other side, RDK based features achieve maximum accuracy of 97.6% and AUC of 0.976 on the training data and validation data using RF-based classifier. Similarly, KNN based models also show comparable results on both training and validation dataset (Table 6). However, XGB performs quite less with an accuracy of 92.5% and an AUC of 0.925 on validation datasets. Results of SVM, DT, and NB classifiers could not perform well on the training as well as validation dataset, as given in Table 6. The complete results are provided in Supplementary Table S6.

**Table 6:** Performance of machine learning based models developed using RDK(K = 3) features on training and validation dataset.

|  | TRAINING |  |  |  |  | VALIDATION |  |  |  |  |
| --- | --- | --- | --- | --- | --- | --- | --- | --- | --- | --- |
| *Classifier* | *Sensitivity* | *Specificity* | *Accuracy* | *AUC* | *MCC* | *Sensitivity* | *Specificity* | *Accuracy* | *AUC* | *MCC* |
| RF | 0.98 | 0.98 | 0.98 | 0.98 | 0.95 | 0.98 | 0.98 | 0.98 | 0.98 | 0.95 |
| KNN | 0.99 | 0.96 | 0.98 | 0.97 | 0.95 | 0.99 | 0.96 | 0.98 | 0.97 | 0.95 |
| SVM | 0.80 | 0.71 | 0.75 | 0.75 | 0.51 | 0.80 | 0.71 | 0.75 | 0.75 | 0.51 |
| DT | 0.95 | 0.94 | 0.95 | 0.95 | 0.90 | 0.95 | 0.95 | 0.95 | 0.50 | 0.00 |
| NB | 0.74 | 0.53 | 0.64 | 0.64 | 0.28 | 0.74 | 0.54 | 0.64 | 0.64 | 0.28 |
| XGB | 0.94 | 0.91 | 0.93 | 0.93 | 0.85 | 0.94 | 0.91 | 0.93 | 0.93 | 0.85 |

#### Web-Server Implementation

To serve the scientific community, we have developed a web-server (https://webs.iiitd.edu.in/raghava/fluspred/) and standalone package (https://github.com/raghavagps/FluSPred) named “FluSPred” using the best model for each protein and the genome datasets. For the frontend part of the web-server, the structure was developed using HTML5, the styling was done by CSS3 and the logic was made using VanillaJS. The backend part was made using Python and PHP. The web-server is responsive as it is compatible with almost all devices including desktop, laptop, mobile phone, tablets, and iMac. Based on the results, the random forest model for one hot encoding feature showed comparable results but even after dimensionality reduction, the models were computer expensive and time consuming, which is why we did not select this for the webserver. On the other hand, execution of the AAC and the DPC models were efficient and so we selected the DPC models for most of the proteins for incorporating them into the web-server since it showed high results in the evaluation parameters compared to AAC and was faster in feature generation compared to the one-hot encoding of the sequences. For NS2 and PB1F2, we chose the AAC models for random forest as they gave the highest results compared to their DPC results. For the genome sequence dataset, CDK feature for K=3 at threshold = 0.5 was incorporated in the web server for the prediction of human and non-human infectious strains using genomic sequences.

The web server has two major modules: (i) Genome and (ii) Protein. User can choose protein/genome-based models according to the requirement. Where, under the Protein module fifteen different models were included. User can select any models for the prediction of the human infectious strain. A single or multiple input sequences can be given by the users in standard FASTA format. The result comprises the prediction of the human/non-human infectious strain, whether the particular strain would be capable of crossing the species barrier to infect humans or not. As a service to the scientific community, the web server “FluSPred” was made open-source. This web-server can be used for further research related to Influenza A or public health and pandemic surveillance in the long run.

#### Case Study: Evaluation of FluSPred

We conducted a study where our web-server was further validated by using new sequence entries, retrieved on 24.12.2021 from IRD. New entries for HA, NP, and PA-X proteins were available. These new sequences were thoroughly checked for redundancy and those were selected which were not a part of the original dataset that was used in the models. For the HA protein, (*A/swine/Iowa/A02636065/2021, A/teal/Samara/Bolshechernigovsky/2021, A/Texas/01/2021*) strains were used, of subtypes H1N2, H5N1, and H3N2 respectively. These sequences were then run on the HA prediction model, which accurately predicted that if the strains were infectious or non-infectious, with the help of probability scores. The score for non-infectious sequences were very low (0.02 and 0.07 for *A/swine/Iowa/A02636065/2021* and *A/teal/Samara/Bolshechernigovsky/2021 strains respectively*) as compared to the human infectious sequence (0.88). Similarly, the sequence of NP protein of *A/Texas/01/2021* strain having H3N2 subtype was used to authenticate the NP prediction model. The result showed was accurate, where the strain cause infections with a score of 0.57. The same strain with PA-X protein sequence was used to test the respective protein model. That model also showed that the strain causes infections to human with a score of 0.98.

## Discussion

Zoonotic diseases, including novel infectious agents (like viruses, bacteria, fungi, protozoa and pathogens), give rise to approximately 2.4 billion cases of illness and 2.7 million deaths [42–45]. Identifying strains that have the susceptibility of causing a disease emergence and can encourage host permissiveness of zoonotic infections is the need of the hour [46]. With the advancements in technology and high-throughput sequencing, now it is possible to develop forecasting tools and early warning systems in disease outbreak analysis. In the last few years, several attempts have been made by scientists worldwide to make computational methods to predict zoonotic events of different pathogens such as influenza, SARS, MERS, Ebola, rabies [24, 47–50]. In literature numerous genome and protein sequence-based prediction methods are developed for the zoonotic host prediction of influenza A virus. Li and Sun [51] used SVM-based approach for the prediction of zoonotic host of rabies virus, coronavirus, influenza A virus, where they achieve an average accuracy of 84%, 85.67%, and 87% respectively [51]. Christine et al., developed computational models by considering 11 influenza proteins and achieve an accuracy of 96.57% and AUC of 0.98 [21]. *Mock et al* [24] developed prediction models using the genomic data of multiple zoonotic species including influenza A virus, rabies lyssavirus and rotavirus A and achieve an average AUC virus species an AUC between 0.93 and 0.98 for various virus species. VIDHOP uses deep neural network approach for the classification and get maximum AUC of 0.94 on the influenza A dataset [24]. Whereas, the methods are based on the zoonotic host prediction of influenza A virus uses limited amount of protein sequence data and do not provide any web-server facility for the community.

In this study, we have made a methodical attempt to develop computational tool using the 15 proteins and genome sequences for the prediction of human infectious strains of the Influenza A virus. The sequence datasets were obtained from IRD and VIDHOP, on which, we calculate compositional-based descriptors/features and one hot encoding-based features using Pfeature and Nfeature tools. Our models included a wide range of data, with 308632 genome sequences pertaining to 34 hosts as well as achieved a much higher accuracy in comparison with existing methods. The relevant features were selected on which the model was trained and validated. Our compositional analysis indicates which residues (A, I, K, E, H, G, P, Q) composition is higher in the positive datasets of three major zoonotic proteins such as HA, NA and PB1-F2. This shows that simple composition-based techniques can identify the important features and help in the prediction. Motif analysis on the sequence data highlighted the important set of motifs that were present on the human sequences and not on the non-human sequences.

The results of this analysis have been provided for use in further research. Here, we used AAC and DPC features for the infectious strain prediction between human (positive) and mammalian /avian(negative) datasets. The AUC varies from 0.88 to 0.99 on validation datasets for 15 different influenza A proteins, with minimum performer PA-N182 and maximum for HA protein. Our results show that Random forest models gave the best prediction, where HA protein achieved the highest AUC of 0.99 in training and 0.98 in validation datasets. Additionally, both HA and NA showed high results on the computational prediction models, which clearly indicate their importance in zoonosis. In case of the genome sequences, random forest showed the best results using CDK (K = 3) features, with an accuracy of 98% and AUC of 0.98 on validation dataset. Of Note, we have used our best models for 15 proteins and genome to implement the web-server named “FluSPred” (https://webs.iiitd.edu.in/raghava/fluspred/). To the best of our knowledge, this is the first attempt to develop a web-server for the prediction of the zoonotic host, whether the particular strain would be capable of crossing the species barrier to infect humans or not, using 15 Influenza A proteins and genome sequence data. The models will be able to predict whether the sequence pertaining to the virus is infectious or not, from avian/mammalian to humans.

## Abbreviations

AUC: Area under the ROC Curve
MCC: Matthews Correlation Coefficient
IRD: Influenza Resource Database
XGB: eXtreme Gradient Boosting
KNN: K-Nearest Neighbor 0 SVM – Support Vector Machine
GNB: Gaussian Naive Bayes
DT: Decision Tree
RF: Random Forest
AAC: Amino Acid Composition
DPC: Dipeptide Composition
CDK: k-mer Composition
RDK: Reverse Compliment k-mer composition

## Authors’ contributions

TR and KS collected and processed the datasets. TR, KS, AD, SP and GPSR implemented the algorithms and developed the prediction models. TR, KS, AD, SP and GPSR analysed the results. SP, KS created the back-end of the web server the front-end user interface. TR, KS, AD, SP and GPSR penned the manuscript. GPSR conceived and coordinated the project. All authors have read and approved the final manuscript.

## Acknowledgements

Authors are thankful to the Department of Bio-Technology (DBT) and Department of Science and Technology (DST-INSPIRE) for fellowships and the financial support and Department of Computational Biology, IIITD New Delhi for infrastructure and facilities.

## Data Availability Statement

All the datasets generated in this study are available at “FluSPred” web server, https://webs.iiitd.edu.in/raghava/fluspred/download.php.

